# From micro-architecture to networks: canonical neurobiological architecture of functional brain organization across the lifespan and in psychopathology

**DOI:** 10.64898/2026.10.22.761188

**Authors:** Guozheng Feng, Jiayu Chen, Armin Iraji, Mohammad S.E. Sendi, Zening Fu, Vince D. Calhoun

## Abstract

Understanding how macroscale functional network connectivity (FNC) across the lifespan and in psychopathology is spatially organized with respect to microscale neurobiological architecture remains a challenge. Here, we constructed a cross-omics mapping framework by linking resting-state fMRI data (5,349 individuals, 6–80 years) with 49 neurobiological markers (24 cell types, 19 neurotransmitters, 6 mitochondria). We show that FNC development and aging unfold along cellular and molecular gradients. Specifically, spatial correspondence between age-related FNC changes and neurobiological microprofile networks varied by life stage, peaking for cell types in adolescence, neurotransmitter systems in mid-young adulthood and mitochondrial phenotypes in early adulthood. Applying normative modeling to 5 psychiatric disorders (schizophrenia, bipolar disorder, major depressive disorder, autism, and attention-deficit/hyperactivity disorder), we identified disorder-specific and cross-disorder FNC phenotypes spatially associated with neurobiological architecture. Together, these findings provide a hypothesis-generating framework linking functional brain organization and dysfunction to canonical neurobiological architecture across the lifespan and psychiatric disorders.

## Introduction

The human brain is a multiscale system whose macroscopic functional organization is embedded within a complex hierarchy of microscale biological substrates^1–4^. A central challenge in neuroscience is how these diverse molecular and cellular features are spatially related to large-scale functional network connectivity (FNC), particularly as FNC varies across the lifespan and deviates in psychopathology. While transcriptomic, cellular, and neuromodulatory atlases have each separately advanced our understanding of brain organization^5–9^, a critical gap remains: a unified, cross-omics framework for systematically testing the alignment between canonical microscale biological architecture and functional connectivity phenotypes.

Normative modeling of large-scale neuroimaging data has gained considerable attention as a powerful approach to capture dynamic brain changes across the lifespan^10–12^. Leveraging large, multi-site cohorts, normative trajectories of FNC across the lifespan have been established, revealing predictable patterns of network maturation and aging^13,14^, and identifying critical inflection points in the development of the connectome^13^. A growing body of evidence indicates that deviations from such normative network patterns play a key role in understanding the pathophysiology of psychiatric disorders^15–17^, helping to reveal both disorder-specific and transdiagnostic alterations in large-scale network connectivity.

Concurrently, the growing availability of high-resolution biological data—from microarray-based transcriptomics, single-nucleus RNA sequencing (snRNA-seq), to positron emission tomography (PET) imaging of neurotransmitter systems—has provided unprecedented opportunities to annotate the brain’s canonical microscale architecture as observed in the general adult population^5–9,18,19^. In neurotypical populations, the spatial distributions of specific cell types and neurotransmitters have been shown to relate to resting-state functional MRI (rs-fMRI) signal amplitude^20^, functional connectivity strengths^21^, principal functional gradient^7^, and intrinsic connectivity networks (ICNs)^22–24^. Beyond spatial correspondence, network-level patterns of cell types, neurotransmitters, and mitochondrial phenotypes exhibit significant correspondence with functional connectivity, suggesting that functionally coupled regions tend to share similar microscale biological profiles^8,22^. Studies have also used neurobiological atlases to contextualize disorder-related brain alterations^20,25–28^. For example, polygenic risk for schizophrenia (SCZ) predicts reduced functional signal amplitude, which in turn negatively correlates with cortical expression of parvalbumin-positive interneurons^20^. Moreover, an analysis across 13 psychiatric conditions showed that regional neurotransmitter receptor profiles were among the strongest predictors of both disorder-specific cortical morphology and cross-disorder similarity^27^. Collectively, these findings support the use of canonical biological atlases as spatial references for annotating macroscale functional and clinical brain phenotypes.

While normative FNC trajectories and deviations from these trajectories across psychiatric disorders have increasingly been established, a systematic, network-level framework for testing whether FNC patterns are organized within specific microarchitectural contexts is still lacking. We hypothesized that normative FNC organization and disorder-related FNC deviations are aligned with canonical microscale biological architecture. To test this hypothesis, we integrated multiscale biological data—encompassing 24 transcriptomically imputed cell-type distributions^5,6^, 19 neurotransmitter receptor/transporter densities^8^, and 6 mitochondrial phenotypic maps^9^—to construct a high-resolution ensemble of neurobiological “microprofile” networks. Distinct from previous studies^8,22^, we computed a network for each biological marker based on its regional probability distributions^29,30^, thereby quantifying the similarity of microscale biological profiles between functional regions. This approach provided a scalable and interpretable way to estimate global and marker-specific correspondence between microprofile networks and FNC phenotypes across modalities. We then related these microprofile networks to lifespan trajectories of FNC derived from a large normative cohort (ages 6–80 years) comprising 3,834 healthy individuals, and estimated FNC deviations of 1,515 patients across five psychiatric disorders: SCZ, bipolar disorder (BP), major depressive disorder (MDD), autism spectrum disorder (ASD), and attention-deficit/hyperactivity disorder (ADHD).

We first examined whether normative FNC patterns at different life stages were differentially associated with distinct types of neurobiological microprofile networks. We then quantified age-related annualized changes in FNC, revealing stage-specific associations between patterns of functional reorganization and corresponding neurobiological architectures. Finally, we applied the normative model to individuals with psychiatric disorders to identify disorder-related deviations in FNC and to assess how these deviations, as well as inter-disorder differences, are aligned with neurobiological microprofile networks. By bridging transcriptomic, cellular, molecular, and systems-level functional data, this work establishes a biologically grounded framework for linking macroscale functional connectivity phenotypes to canonical microscale biological architecture across the lifespan and in psychopathology.

## Results

We constructed a large-scale, multi-site dataset of FNC using the Neuromark framework with functional 1.0 template^31^, comprising a total of 5,349 participants drawn from both lifespan cohorts (HCP-D^32^, HCP-YA^33^, and HCP-A^34^) and datasets covering five psychiatric disorders: SCZ (FBIRN^35^ and COBRE^36^), BP (LA5C^37^), MDD (Chinese Han MDD [CMDD]^38^), ASD (ABIDE I and II^39,40^), and ADHD (ADHD200^41^). Detailed demographic information is provided in ***Supplementary Table 1***. Using all neurotypical participants in these datasets, we built a normative model to derive multiple FNC phenotypes, including normative connectivity patterns, their age-related changes across the lifespan, and disorder-related alterations in each of the five psychiatric conditions. To assess the neurobiological associations, we systematically examined the connectional correspondence between these FNC phenotypes and a diverse set of neurobiological microprofile networks derived from cell types^6^, neurotransmitter systems^8^, and mitochondrial phenotypes^9^. The overall framework is illustrated in **Fig. 1**.

**Fig. 1.**
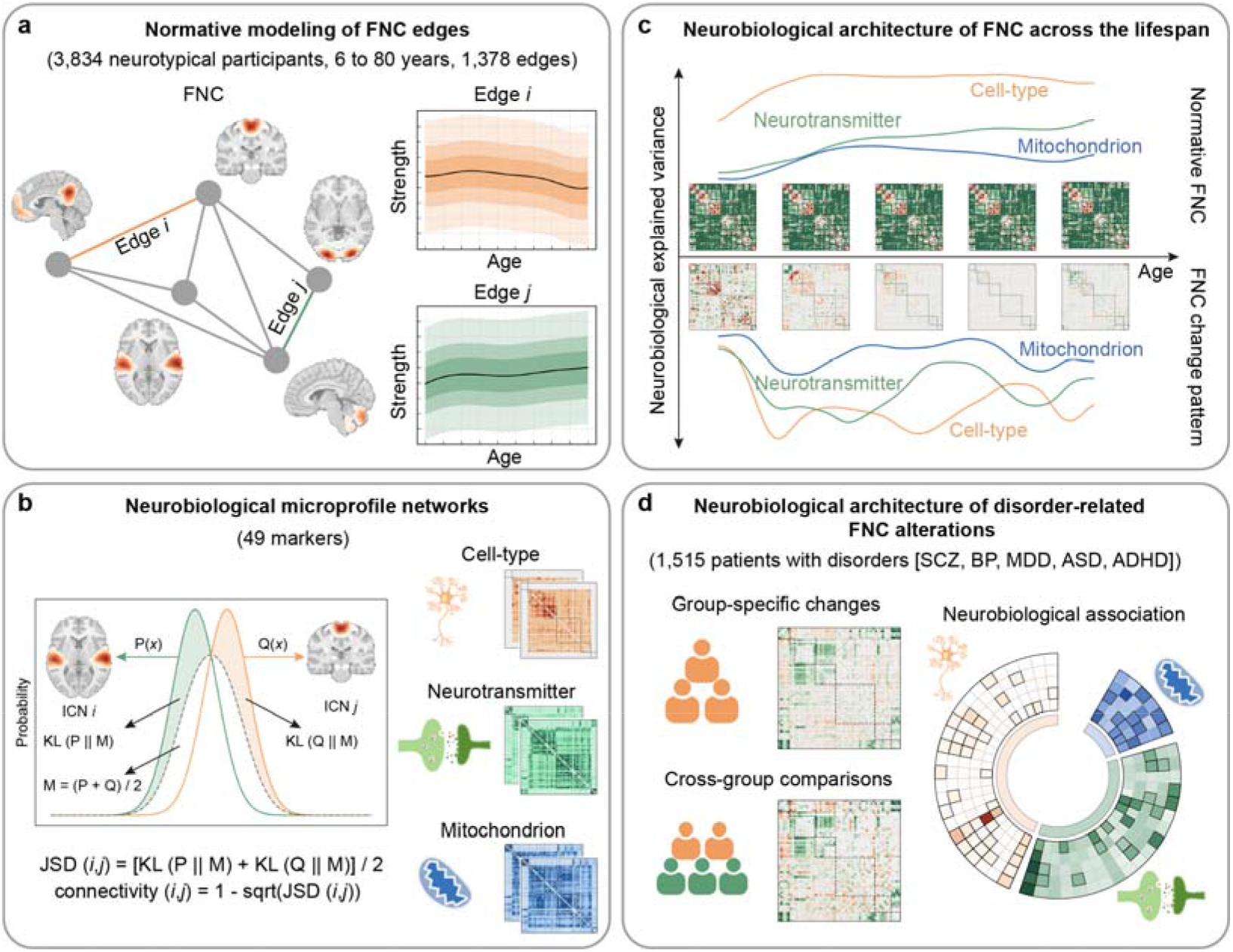
| Overview of the analytical framework. **a**, Normative modeling of 1,378 FNC edges, defined by the Neuromark framework with functional 1.0 template^31^, was performed using the PCNtoolkit^43^ based on data from 3,834 healthy participants (aged 6–80 years). **b**, Neurobiological microprofile networks were constructed using the spatial distributions of 49 biological markers, including 24 imputed cortical cell types^5,6^, 19 neurotransmitter systems^8^, and 6 mitochondrial phenotypes^9^. Pairwise similarity between ICNs was calculated using Jensen–Shannon divergence (JSD), with similarity defined as 1−sqrt(JSD). **c**, Normative FNC expectations across the lifespan (ages 6–80) were estimated. FNC change patterns were computed as one-year differences between consecutive normative FNC estimates (i.e., FNC at age *t* minus FNC at age *t*−1). The spatial correspondence of normative FNC patterns and age-related FNC change patterns with neurobiological microprofile networks was assessed. **d**, Individual FNC deviations (d-scores) were computed by applying the normative model to 1,515 patients with disorders (five groups: SCZ, BP, MDD, ASD, ADHD), and group-specific and cross-group difference maps were obtained. The spatial correspondence of disorder-specific and cross-disorder maps with neurobiological microprofile networks was then assessed.

### Alignment between normative FNC across the lifespan and canonical neurobiological architecture

We constructed a normative model of FNC across the lifespan using Bayesian linear regression (BLR)^42^, as implemented in the Predictive Clinical Neuroscience Toolkit (PCNtoolkit)^43^. The model was trained on FNC data from 3,834 neurotypical participants across 1,378 connectivity edges. Normative FNC trajectories were estimated at 75 integer age points spanning ages 6 to 80, covering five major life stages: late childhood (6–12 years), adolescence (12–20 years), young adulthood (20–40 years), middle adulthood (40–60 years), and late adulthood (60+ years)^44^ (***Supplementary Fig. 1***). To assess whether these normative FNC patterns are aligned with canonical neurobiological architecture, we integrated 49 neurobiological marker images, encompassing 24 cell types^6^, 19 neurotransmitter receptors and transporters^8^, and 6 mitochondrial phenotypes^9^. For each marker image, we computed pairwise regional similarity using Jensen–Shannon divergence (JSD)^29,30^, resulting in 49 marker-specific neurobiological microprofile networks aligned with FNC (***Supplementary Fig. 2***). The abbreviations and full names of the neurobiological markers are detailed in ***Supplementary Table 2***. We assessed the biological architecture of normative FNC by modeling its correspondence with the 49 neurobiological microprofile networks. For each age, we separately fit multivariate linear models^45,46^ at three biological scales—cell-type, neurotransmitter, and mitochondrial microprofile networks—with the normative FNC pattern as the dependent variable. We then evaluated both the overall explained variance (adjusted *r*²) and the relative contribution of individual markers within each scale (see **Methods** for full details).

Across the lifespan, multivariate analyses revealed sustained colocalization between FNCs and neurobiological microprofile networks across life stages for all three categories: cell types (adjusted *r*^2^ = 0.21 ± 0.01, **Fig. 2a**), neurotransmitters (adjusted *r*^2^ = 0.14 ± 0.02, **Fig. 2a**), and mitochondria (adjusted *r*^2^ = 0.12 ± 0.01, **Fig. 2a**). All associations were statistically significant (false discovery rate [FDR] *q* < 0.05). Relative importance analysis further revealed marker-specific contributions throughout the lifespan (**Fig. 2b**). Among cell-type microprofile networks, layer 6b neurons (L6b), astrocytes (Astro), oligodendrocyte precursor cells (OPC), and oligodendrocytes (Oligo) consistently exhibited high relative importance across life stages, with Astro emerging as the top contributor at nearly all ages. For neurotransmitter systems, serotonin 1A receptor (5-HT1a), norepinephrine transporter (NET), and cannabinoid CB1 receptor (CB1) showed persistent contributions across the lifespan, with NET being the most dominant across nearly all stages. Among mitochondrial markers, complex I (CI) and mitochondrial density (MitoD) robustly explained FNC variance throughout the lifespan. The univariate associations showed consistent patterns (***Extended Data Figs. 1–3***).

**Fig. 2.**
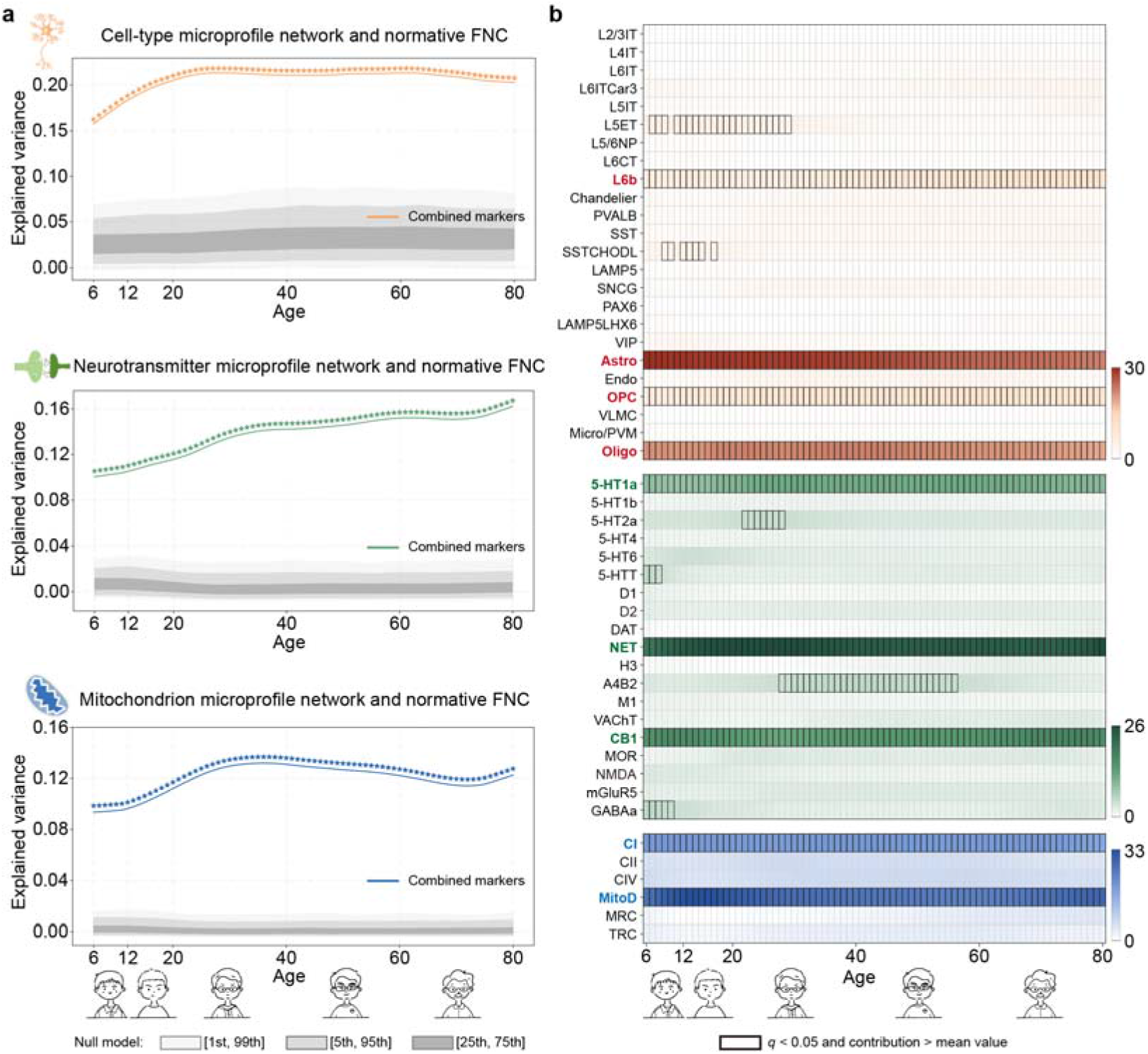
| Colocalization between cross-sectional normative FNC and neurobiological microprofile networks across the lifespan. **a**, Normative FNC expectations were sampled at 75 integer age points across the lifespan. At each age, spatial correspondence was assessed between the cross-sectional FNC and neurobiological microprofile networks derived from cell types, neurotransmitter systems, and mitochondrial phenotypes, using multiple linear regression models. Spatial confounds (Euclidean distance and Dice similarity) were included as covariates and regressed out. The adjusted *r*² was used to quantify the proportion of FNC variance explained by each model. Statistical significance was evaluated via one-sided permutation testing against 1,000 FNC surrogate maps, with FDR correction applied across 75 age-point comparisons. Smoothed regression curves were fitted using locally estimated scatterplot smoothing (LOESS) across the observed explained variance and the null distribution percentiles (1st, 5th, 25th, 75th, 95th, 99th) to highlight developmental trajectories. Colored asterisks along the curves indicate age points with significant model performance after permutation testing and FDR correction (*q* < 0.05). **b**, Relative importance analysis^45,46^ was performed to partition the relative contribution across predictors. Statistical significance was assessed using one-sided permutation testing against 1,000 FNC surrogate maps, with FDR correction applied across 75 × (24, 19, or 6) comparisons. Predictors that were statistically significant and showed a relative contribution exceeding the average expected contribution (i.e., 1/(24, 19, or 6)) were considered to exhibit high explanatory influence.

### Alignment between FNC changes across the lifespan and canonical neurobiological architecture

Having established widespread colocalization between cross-sectional normative FNC and neurobiological microprofile networks, we next investigated whether FNC-change patterns are aligned with canonical microprofile architectures. We computed annualized FNC change at each of the 74 integer age points (7–80 years) as differences between consecutive normative FNC estimates (e.g., FNC at age *t* minus FNC at age *t*−1), capturing dynamic reorganization over time. For each age, we assessed the relationship between FNC change and the three microprofile modalities using separate multivariate linear models^45,46^ (see **Methods**).

Multivariate models confirmed modality-specific explanatory power (**Fig. 3a**): cell types (adjusted *r*^2^ = 0.10 ± 0.03), neurotransmitters (adjusted *r*^2^ = 0.08 ± 0.03), and mitochondria (adjusted *r*^2^ = 0.03 ± 0.02). All three models showed statistically significant variance explained at nearly all evaluated ages (*q* < 0.05). The peak explanatory power of cell-type features was observed during adolescence, whereas neurotransmitter systems peaked in mid–young adulthood and mitochondrial contributions reached their maximum in early adulthood (**Fig. 3a**). Relative importance analysis revealed stage-specific contributions of individual neurobiological markers (**Fig. 3b**). Several markers exhibited pronounced age-specific trajectories, with maximal contributions around major developmental transitions. For example, layer 6 intratelencephalic neurons (L6IT) showed preferential contributions from young adulthood through late adulthood, with a peak in early middle adulthood. The correspondence of Oligo and 5-HT1a microprofile networks with FNC-change patterns increased from childhood to adolescence and decreased thereafter. CB1 showed particularly high contributions around the transition from middle to late adulthood, whereas mitochondrial respiratory capacity (MRC) exhibited preferentially high contributions around the transition from young to middle adulthood. The univariate associations showed consistent patterns (***Extended Data Figs. 4–6***).

**Fig. 3.**
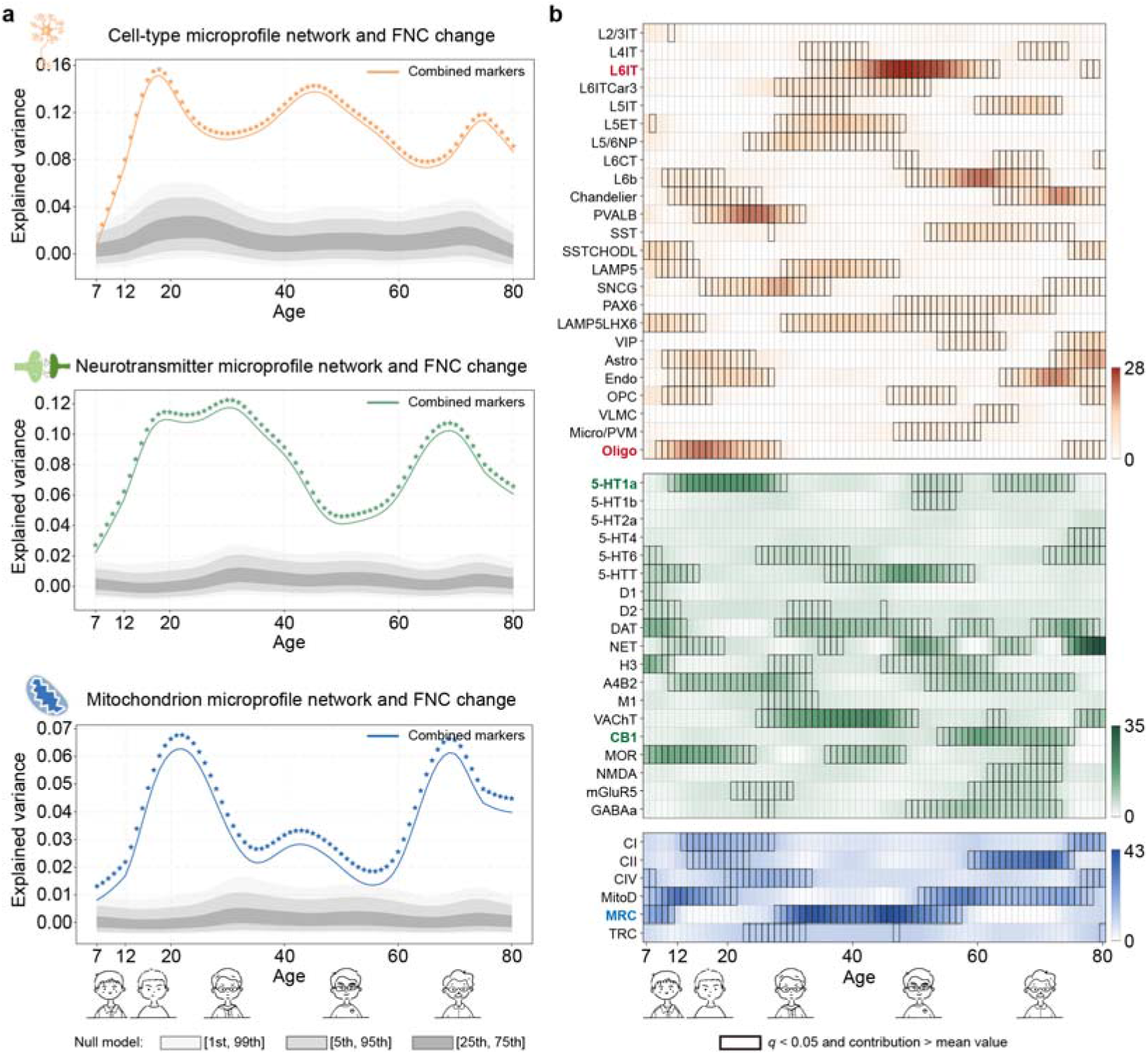
| Modeled FNC change patterns explained by neurobiological microprofile networks. **a**, Age-related changes in FNC were estimated as differences between consecutive normative FNC expectations, yielding 74 annualized change maps for ages 7–80 years. These annualized change maps were then associated with neurobiological microprofile networks derived from cell types, neurotransmitter systems, and mitochondrial phenotypes, using multiple linear regression models. Spatial confounds (Euclidean distance and Dice similarity) were included as covariates and regressed out. The adjusted *r*² was used to quantify the proportion of variance in FNC changes explained by each model. Statistical significance was evaluated via one-sided permutation testing against 1,000 surrogate maps, with FDR correction applied across 74 age-point comparisons. Smoothed regression curves were fitted using LOESS across the observed explained variance and the null distribution percentiles (1st, 5th, 25th, 75th, 95th, 99th) to highlight developmental trajectories. Colored asterisks along the curves indicate age points with significant model performance after permutation testing and FDR correction (*q* < 0.05). **b**, Relative importance analysis^45,46^ was performed to partition the relative contribution across predictors. Statistical significance was assessed using one-sided permutation testing against 1,000 FNC surrogate maps, with FDR correction applied across 74 × (24, 19, or 6) comparisons. Predictors that were statistically significant and showed a relative contribution exceeding the average expected contribution (i.e., 1/(24, 19, or 6)) were considered to exhibit high explanatory influence.

Our analyses aimed to characterize the spatial correspondence between lifespan-related FNC patterns and canonical neurobiological architecture derived from static reference atlases, rather than to infer age-dependent changes in the underlying biological features. To assess the suitability of these atlases as spatial references, we performed two complementary validation analyses. Allen Human Brain Atlas (AHBA) donor-level leave-one-donor-out analysis revealed moderate reproducibility of cell-type microprofile networks (mean Spearman’s *r* = 0.42 ± 0.18; ***Extended Data Fig. 7***). An independent lifespan transcriptomic dataset^44^ spanning ages 8–70 years, selected using an age inclusion criterion of 6–80 years, showed no significant age-related drift in regional expression contrast patterns across the 24 cell types (***Extended Data Fig. 8***).

### Disorder-specific FNC deviations align with canonical neurobiological architecture

Using the normative model, we computed individual FNC deviation scores (d-scores) for each patient, quantifying subject-specific departures from expected normative connectivity. A total of 1,515 patients were included across five diagnostic groups: SCZ (*n* = 219), BP (*n* = 46), MDD (*n* = 256), ASD (*n* = 764), and ADHD (*n* = 230). Disorder-specific alterations were identified using a two-sided one-sample *t*-test on edge-wise d-scores (**Fig. 4a**; FDR *q* < 0.05). We observed distinct alteration patterns across disorders. For example, SCZ exhibited the most widespread deviations, characterized by decreased connectivity between sensorimotor and visual domains (SM–VIS) and between subcortical and cerebellar domains (SC–CB), alongside increased connectivity between subcortical and auditory domains (SC–AUD) and between cognitive control and default mode domains (CC–DM).

**Fig. 4.**
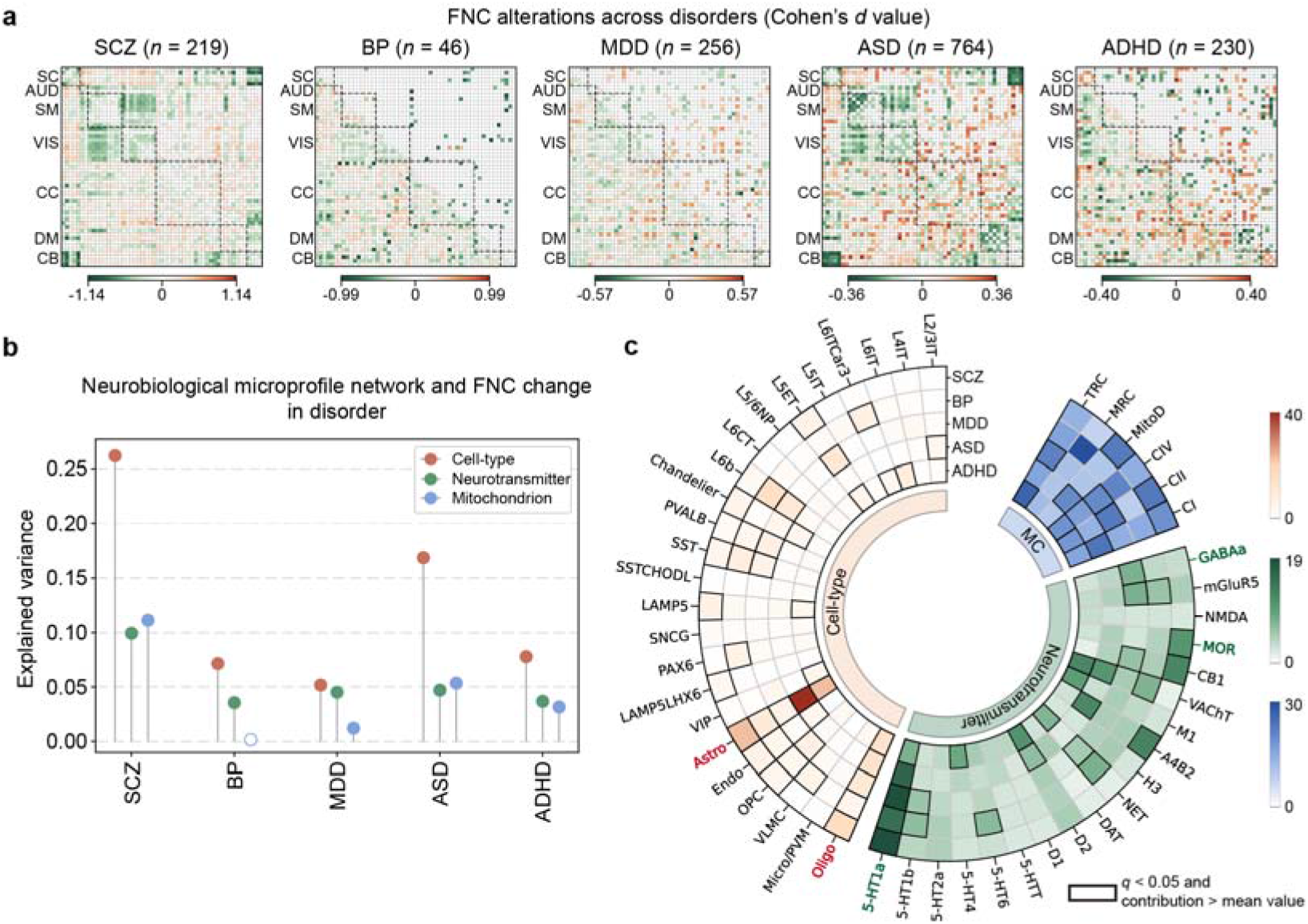
| Disorder-specific FNC alterations and associations with neurobiological microprofile networks. **a**, Disorder-specific FNC alterations are presented as Cohen’s *d* maps (lower triangle). Two-sided one-sample *t*-tests were used to identify connections that significantly differed from zero (upper triangle), with FDR correction applied across 1,378 comparisons. **b**, Disorder-specific Cohen’s *d* maps were then associated with neurobiological microprofile networks derived from cell types, neurotransmitter systems, and mitochondrial phenotypes using multiple linear regression. Spatial confounds (Euclidean distance and Dice similarity) were included as covariates and regressed out. The adjusted *r*² quantified the proportion of variance explained by each model. Statistical significance was assessed via one-sided permutation testing against 1,000 surrogate maps, with FDR correction across 5 comparisons. Filled circles indicate significant associations after FDR correction, whereas open circles indicate non-significant associations. **c**, Relative importance analysis^45,46^ was performed to partition the relative contribution across predictors. Statistical significance was evaluated using one-sided permutation testing against 1,000 surrogate maps, with FDR correction applied across 5 × (24, 19, or 6) comparisons. Predictors that were significant and exhibited a relative contribution exceeding the average expected contribution (i.e., 1/(24, 19, or 6)) were considered to have high explanatory influence. Color intensity indicates relative importance (%), and black outlines indicate predictors meeting both the FDR significance and above-average contribution criteria.

To relate these disorder-specific deviations to canonical neurobiological architecture, we separately modeled the correspondence between each disorder’s Cohen’s *d* map and the cell-type, neurotransmitter, and mitochondrial microprofile networks and assessed both overall explained variance and marker-level contributions (see **Methods**). Across disorders, cell-type and neurotransmitter features consistently explained significant variance in FNC deviations (**Fig. 4b**; FDR *q* < 0.05), and mitochondrial features were significant for all disorders except BP. SCZ showed the strongest neurobiological correspondence across modalities (cell-type adjusted *r*² = 0.26; neurotransmitter = 0.10; mitochondrion = 0.11). Relative importance analysis further revealed both disorder-specific and cross-disorder marker-level contributions (**Fig. 4c**). For example, the mu-opioid receptor (MOR) was selectively dominant in SCZ, whereas gamma-aminobutyric acid type A receptor (GABAa) showed specific contributions in MDD. Conversely, Astro, Oligo, and 5-HT1a exhibited dominant contributions across multiple disorders, suggesting that disorder-related FNC deviations may be organized along shared canonical neurobiological similarity axes. Univariate analyses showed consistent patterns (***Extended Data Fig. 9***).

### Cross-disorder differences in FNC alterations align with canonical neurobiological architecture

To characterize differences in FNC alteration patterns across disorders, we quantified pairwise group differences in d-score maps using a two-sided two-sample *t*-test. All ten contrasts (Cohen’s *d*) revealed substantial numbers of significantly different connections (FDR *q* < 0.05, **Fig. 5a**), indicating robust disorder-specific divergence in FNC alteration patterns. We next examined whether these cross-disorder differences were aligned with canonical neurobiological architecture. For each contrast, we modeled its correspondence with the cell-type, neurotransmitter, and mitochondrial microprofile networks (see **Methods**). Multiple neurobiological scales significantly accounted for cross-disorder FNC differences (**Fig. 5b**; FDR *q* < 0.05), suggesting that differences between disorder-related FNC deviations are organized along distinct canonical biological similarity axes. Relative importance analysis further revealed both disorder-pair–specific and transdiagnostic marker contributions (**Fig. 5c**). Several markers selectively characterized specific contrasts—for example, Oligo and Chandelier were particularly dominant in comparisons involving SCZ and other disorders, while the MOR differentiated SCZ and ADHD from the remaining groups. Notably, 5-HT1a contributed significantly to most cross-disorder contrasts, suggesting that cross-disorder FNC differences may be organized along a broadly shared canonical serotonergic similarity axis. Univariate analyses showed consistent patterns (***Extended Data Fig. 10***).

**Fig. 5.**
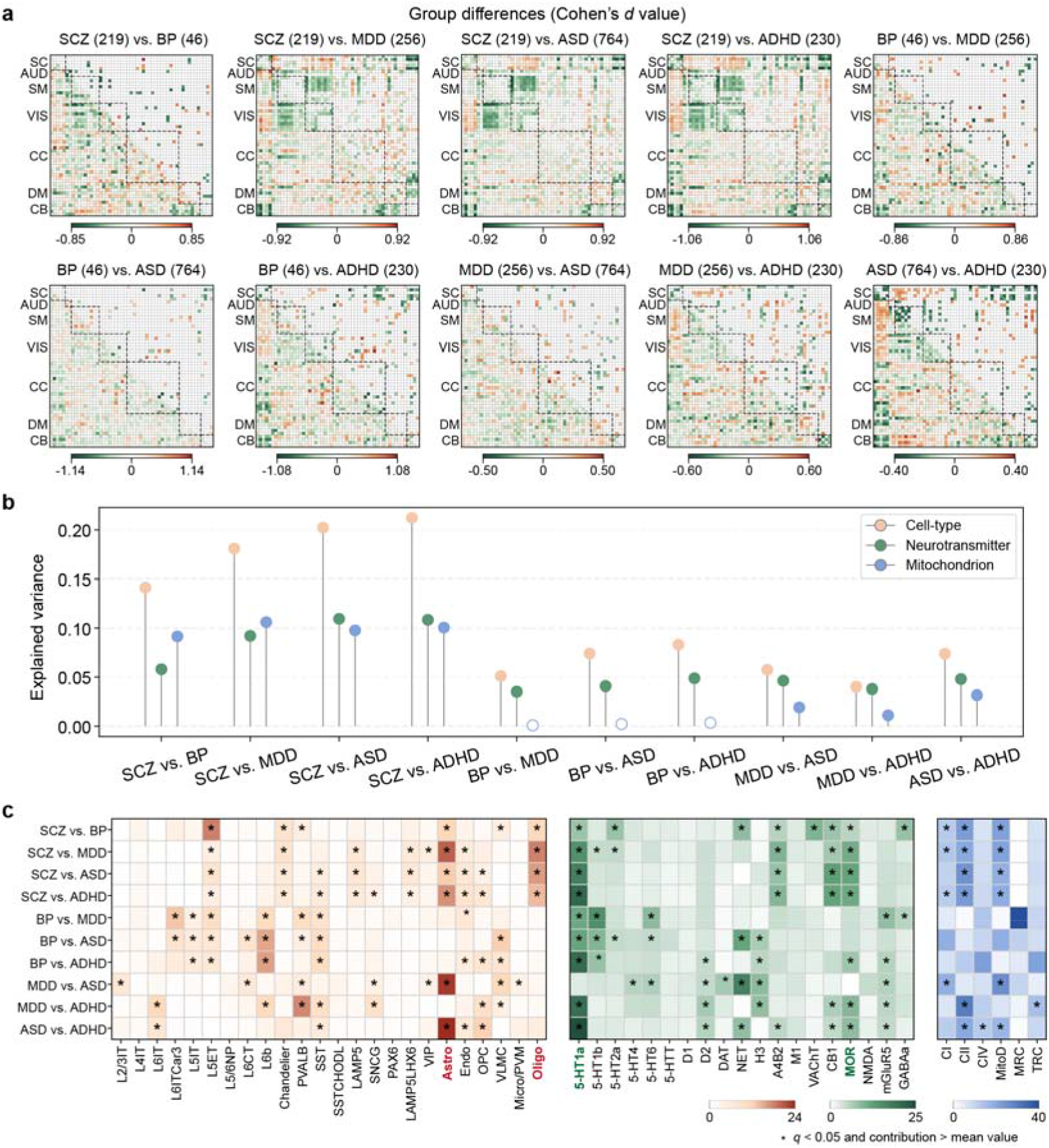
| Cross-disorder differences in FNC alterations and associations with neurobiological microprofile networks. **a**, Group differences in FNC d-score maps between disorder pairs were assessed using two-sided two-sample *t*-tests, with effect sizes quantified by Cohen’s *d* (lower triangle). FDR correction was applied across 1,378 comparisons to identify connections showing significant cross-disorder differences. Numbers in parentheses indicate the number of patients in each diagnostic group. **b**, These cross-disorder difference maps were then associated with neurobiological microprofile networks derived from cell types, neurotransmitter systems, and mitochondrial phenotypes using multiple linear regression. Spatial confounds (Euclidean distance and Dice similarity) were included as covariates and regressed out. The adjusted *r*² quantified the proportion of variance in cross-disorder d-score differences explained by each neurobiological model. Statistical significance was determined via one-sided permutation testing against 1,000 surrogate maps, with FDR correction applied across 10 comparisons. Filled circles indicate significant associations after FDR correction, whereas open circles indicate non-significant associations. **c**, Relative importance analysis^45,46^ was performed to partition the relative contribution across predictors. Statistical significance was assessed using one-sided permutation testing against 1,000 surrogate maps, with FDR correction applied across 10 × (24, 19, or 6) comparisons. Predictors that were statistically significant and exhibited a relative contribution exceeding the average expected contribution (i.e., 1/(24, 19, or 6)) were considered to have high explanatory influence. Color intensity indicates relative importance (%), and black outlines indicate predictors meeting both the FDR significance and above-average contribution criteria.

## Discussion

This study directly addressed a key gap by establishing a coherent, multiscale bridge relating macroscale functional connectivity phenotypes across the lifespan and in psychiatric disorders to canonical microscale biological architecture. By integrating normative modeling of FNC with a comprehensive ensemble of neurobiological microprofile networks, we found that normative FNC organization, age-related FNC-change patterns, and disorder-related FNC deviations are spatially organized along canonical microscale biological axes, with the strength and marker-specific pattern of this correspondence varying across life stages and psychiatric conditions.

Normative modeling of large-scale neuroimaging data has gained considerable attention as a powerful approach to capture dynamic brain changes across the lifespan and deviations associated with neuropsychiatric disorders^10,11,13,47–49^. Spatial colocalization with canonical microscale biological atlases can provide an exploratory framework for annotating the biological architecture associated with both normative and atypical human brain processes. However, because these canonical atlases are typically static, group-level, and often adult-derived, and in some cases based on postmortem samples, inferences from spatial colocalization analyses are generally limited to the associative rather than causal level. Our additional validation analyses indicated moderate reproducibility of cell-type microprofile similarity across AHBA donors. An independent lifespan transcriptomic analysis found no significant age-related drift in regional expression contrast patterns across the 24 cell types. These findings do not imply that the biological atlases capture age-specific cellular or molecular changes. Rather, they provide qualified support for using canonical microscale biological architecture as a spatial reference for examining the correspondence with lifespan-related and disorder-related FNC phenotypes.

Previous work has shown that normative trajectories of regional cortical thickness can be partially explained by spatial distributions of neurobiological markers, with distinct markers contributing at different developmental stages^50^. Extending this framework to FNC, we revealed stable, network-level colocalization between lifespan normative FNC patterns and neurobiological microprofile networks, supporting the idea that functionally coupled regions may tend to share similar microscale biological profiles^51^. These shared biological profiles may facilitate participation in similar neural computations and coordinated network dynamics. Although the observed associations are broadly consistent with prior multidisciplinary evidence, such convergence should be interpreted as supportive biological plausibility rather than proving that our findings reflect true development-related neurobiological mechanisms. Several markers, including astrocytes, oligodendrocytes, NET, and CB1, showed sustained high relative importance for functional architecture across the lifespan. These markers have established roles in processes relevant to functional coordination: astrocytes regulate neuronal rhythmicity, synchrony, and global brain states^52^; oligodendrocytes influence conduction velocity and network synchrony through activity-dependent myelination^53^; NET regulates noradrenergic signal clearance and exhibits spatial correspondence with brainstem–cortical functional organization^54,55^; and CB1 receptors modulate synaptic transmission and neural oscillatory synchrony^56,57^. We further found that the correspondence between FNC-change patterns and microprofile networks varied across age, with cell-type networks showing the strongest correspondence during adolescence and neurotransmitter networks showing the strongest correspondence in young adulthood. These age-dependent patterns parallel observations from normative cortical thickness trajectories^50^ and are broadly consistent with prior evidence that myelination undergoes pronounced refinement during adolescence and continues into young adulthood^58–60^, and that the default mode network—a late-developing system modulated by 5-HT1a receptors^61,62^—continues to mature throughout adolescence and early adulthood.

Psychiatric disorders are associated with multilevel alterations spanning molecular and cellular processes, neural dynamics, and large-scale functional networks. Recent studies have used neurobiological atlases to contextualize disorder-related brain changes and identify biological axes along which macroscale abnormalities may be spatially organized^20,25,27,28,63^. In this study, we quantified both disorder-specific FNC d-scores and inter-disorder differences, revealing statistically robust alterations that parallel prior reports of a spectrum of network-level disruptions across psychiatric conditions^64^. By aligning these disorder-related FNC phenotypes with canonical neurobiological microprofile networks, we found that astrocytes, oligodendrocytes, and 5-HT1a receptors showed consistently high relative importance across multiple disorders, suggesting that disorder-related FNC deviations may be organized along shared canonical glial and neuromodulatory similarity axes. This interpretation is broadly consistent with prior evidence implicating astrocytes, oligodendrocytes, and serotonergic systems in processes relevant to large-scale functional organization. Astrocyte dysfunction has been linked to brain-wide abnormalities in functional connectivity and cognition^65^, contributes to SCZ^66^, and extends to mood and neurodevelopmental disorders, such as MDD^65^ and ASD^67^; likewise, oligodendrocyte and myelin dysfunction has been repeatedly observed in SCZ^68^ and MDD^69^. Serotonin plays a central role in regulating plasticity, cortical circuit development, and large-scale network dynamics^62^, and 5-HT1a receptors have been broadly implicated in major neuropsychiatric conditions and represent important therapeutic targets^70^. Together, these findings provide hypothesis-generating biological axes for future studies using patient-derived molecular data, longitudinal imaging, or pharmacological intervention designs^50,71,72^.

While our findings provide network-level insights into the neurobiological foundations of FNC across both normative development and psychiatric disorders, several limitations should be acknowledged. First, due to the absence of non-invasive tools for developmental neurobiology in humans, our inferences rely on spatial colocalization analyses, which are inherently associative and non-causal. As such, all reported associations should be interpreted as indirect evidence of involvement, rather than mechanistic proof of specific neurobiological processes. Second, the cellular and molecular maps used in this study were derived primarily from adult neurotypical donor-based atlases. Although these atlases provide valuable canonical references for linking macroscale functional organization to microscale neurobiological architecture, they may not capture inter-individual variability related to age, sex, genetic background, or disease-specific alterations in cellular composition and molecular signaling. Third, although we sought to perform a whole-brain analysis, our cell-type maps were restricted to the cerebral cortex, excluding subcortical structures and the cerebellum. These regions are also underrepresented in certain molecular atlases, which may obscure important contributions from these areas in both normative and pathological conditions. Finally, limited statistical power may have reduced sensitivity to detect additional effects, particularly for BP. Future studies with larger, multi-center cohorts will be important to enhance statistical power, improve generalizability, and further validate the present findings.

These findings provide a scalable cross-omics framework for biologically annotating functional network alterations across the lifespan and in psychopathology. By mapping FNC phenotypes onto canonical cellular, molecular, and metabolic architectures, this work offers a hypothesis-generating roadmap for future studies aimed at testing the biological mechanisms underlying brain network organization and dysfunction using developmental, patient-derived, longitudinal, or interventional datasets.

## Methods

### Participants

A total of 5,349 participants were included from a combination of lifespan and clinical datasets. Lifespan data were obtained from HCP-D^32^, HCP-YA^33^, and HCP-A^34^. Data for five psychiatric disorders were obtained from FBIRN^35^ and COBRE^36^ (SCZ), LA5C^37^ (BP), CMDD^38^ (MDD), ABIDE I and ABIDE II^39,40^ (ASD), and ADHD200^41^ (ADHD). Of the total sample, 3,834 participants were classified as neurotypical, and 1,515 were diagnosed with one of the five psychiatric disorders. Detailed demographic information was provided in *Supplementary Table 1*. Written informed consent was obtained from participants and/or their legal guardians, and the recruitment procedures were approved by the local ethics committees for each dataset.

### Imaging preprocessing

For each dataset, we used the available preprocessed rs-fMRI data when provided. For datasets lacking preprocessing, we performed standardized preprocessing steps using a combination of the FMRIB Software Library (FSL v6.0, https://fsl.fmrib.ox.ac.uk/fsl/fslwiki/) and Statistical Parametric Mapping (SPM12, http://www.fil.ion.ucl.ac.uk/spm/) toolboxes within the MATLAB 2024b environment.

For FBIRN, COBRE, ABIDE, CMDD, and ADHD200, the first five scans were removed for signal equilibrium and participants’ adaptation to scanner noise. We performed rigid-body motion correction using the toolbox in SPM to correct subject head motion, followed by slice-timing correction to account for differences in slice acquisition timing. The rs-fMRI data were subsequently warped into the standard Montreal Neurological Institute (MNI) space using an echo-planar imaging (EPI) template and were resampled to 3 × 3 × 3 mm^3^ isotropic voxels. The resampled fMRI images were further smoothed using a Gaussian kernel with a full width at half maximum (FWHM) = 6 mm.

For HCP datasets, we discarded ten initial volumes with large signal changes to allow the tissue to reach a steady state of radiofrequency excitation. Second, we corrected head motion using the mcflirt function in FSL and co-registered all the volumes to the first volume. Third, for susceptibility-induced distortion correction, pairs of images acquired with opposite phase-encoding directions (left-to-right/right-to-left for HCP-YA and anterior-to-posterior/posterior-to-anterior for HCP-D and HCP-A) were used with FSL topup^73,74^ to estimate the off-resonance field. The output field map coefficients were used to correct the distortion in the rs-fMRI volume using the FSL tool applytopup. Fourth, after the distortion correction, we performed slice-timing correction within each volume to correct the time difference within each repetition time (TR) using tools incorporated in SPM12. Fifth, rs-fMRI data were warped into the standard MNI space, resampling to 3 × 3 × 3 mm^3^ isotropic voxels using the normalization tool in SPM. Finally, the resliced fMRI images were smoothed using a Gaussian kernel with a FWHM = 6 mm.

### FNC

Preprocessed fMRI data were decomposed into 53 subject-specific independent components and their corresponding time courses (TCs) using a spatially constrained single-subject independent component analysis (ICA) approach guided by the Neuromark functional 1.0 template^31^. The use of 53 components was determined by this template, which comprises ICNs that have been shown to be highly reproducible and biologically meaningful across independent ICA decompositions of multiple large-N cohorts following standard ICN selection criteria. The template is available through the GIFT toolbox^75^ (http://trendscenter.org/software/gift) and at http://trendscenter.org/data. To further reduce residual noise in the ICN TCs, four postprocessing steps were applied: (1) detrending to remove linear, quadratic, and cubic signal trends; (2) outlier detection and removal; (3) multiple regression of six motion parameters (three translational and three rotational) and their temporal derivatives; and (4) temporal band-pass filtering (0.01–0.15 Hz). Pairwise Pearson’s correlations were then computed between all denoised ICN TCs to generate subject-level FNC matrices. To focus on positive functional associations, all negative correlation values were set to zero.

### AHBA microarray data processing

Human brain gene expression data from 3,702 bulk tissue samples across six postmortem adult donors were obtained from the AHBA dataset (http://human.brain-map.org/)^5^. Raw microarray data were processed at the sample level using the abagen toolbox (https://abagen.readthedocs.io/en/stable)^76,77^, following a standardized and widely adopted preprocessing pipeline^76^. Probe reannotation was performed using updated mappings from the reference^76^, and probes lacking valid Entrez Gene IDs were excluded. To ensure data quality, only probes with expression levels exceeding background in at least 30% of tissue samples were retained. For genes with multiple probes, the probe exhibiting the highest differential stability across donors was selected to represent the gene. This procedure yielded expression profiles for a total of 16,383 genes. For normalization, within each donor, gene expression values were first z-scored across genes within each tissue sample, and then z-scored across all samples to mitigate inter-sample variability.

### Cell-type deconvolution

Single-nucleus droplet-based sequencing (snDrop-seq) data^6^ were obtained from the cellxgene-census database (https://cellxgene.cziscience.com/collections/d17249d2-0e6e-4500-abb8-e6c93fa1ac6 f). Gene identifiers were harmonized across datasets by matching Entrez Gene IDs between the snDrop-seq and AHBA datasets, retaining only genes common to both. To reduce computational complexity and enhance biological specificity, we restricted the analysis to reliable marker genes that were consistently detected across eight cortical regions, as identified in reference^6^. To mitigate collinearity in downstream analyses, transcriptionally similar cell types were grouped based on higher-order subclass classifications across cortical areas^6,78^. The aggregated snDrop-seq expression data across the eight regions served as the reference for cell-type deconvolution of the AHBA bulk transcriptomic data, performed using CIBERSORTx (https://cibersortx.stanford.edu/)^79^, a strategy also adopted in similar studies^7,20,22^. This yielded estimated relative abundances for 24 distinct cortical cell types, including L2/3IT, L4IT, L6IT, L6ITCar3, L5IT, L5ET, L5/6NP, L6CT, L6b, Chandelier, PVALB, SST, SSTCHODL, LAMP5, SNCG, PAX6, LAMP5LHX6, VIP, Astro, Endo, OPC, VLMC, Micro/PVM, and Oligo, each with distinct laminar distributions, morphological features, developmental lineages, and projection patterns^6,78^. For each AHBA donor, estimated cell-type abundances were projected into MNI space at a 3 × 3 × 3 mm³ isotropic resolution using the corresponding spatial coordinates of tissue samples. Final group-level spatial distributions for each cell type were derived by averaging across donors^7^.

### Molecular atlas

We included two sets of spatial molecular maps characterizing neurotransmitter systems and mitochondrial phenotypes. Neurotransmitter receptor and transporter density maps were derived from 19 PET-based tracer images obtained from the publicly available repository at https://github.com/netneurolab/hansen_receptors^8^. The analyzed receptors and transporters included 5-HT1a, 5-HT1b, 5-HT2a, 5-HT4, 5-HT6, 5-HTT, D1, D2, DAT, NET, H3, A4B2, M1, VAChT, CB1, MOR, NMDA, mGluR5, and GABAa. In parallel, six mitochondrial phenotype maps were retrieved from https://neurovault.org/collections/16418/^9^, including CI, CII, CIV, MitoD, TRC, and MRC. All molecular images were coregistered to MNI space at a 3 × 3 × 3 mm³ isotropic resolution. This ensured consistent spatial alignment across imaging, transcriptomic, and molecular modalities for subsequent multimodal integration analyses.

### Neurobiological microprofile network

For each neurobiological spatial map spanning cellular and molecular features, we adopted a divergence-based framework, originally developed for morphological network construction^29^, to compute marker-specific microprofile networks by quantifying the pairwise similarity of regional probability distributions. Specifically, we computed the JSD, a symmetric and bounded measure from information theory that quantifies the dissimilarity between two probability distributions^29,30^. The JSD between two distributions P and Q is defined as

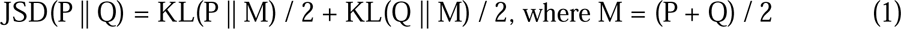

Here, KL denotes the Kullback–Leibler divergence, and M is the average distribution. The resulting JSD is a symmetrized, smoothed, and bounded divergence measure with strong theoretical properties. In our context, P(*x*) and Q(*x*) represent the spatial distributions of a biological marker across two ICNs. Pairwise similarity between brain regions was then defined as 1−sqrt(JSD), resulting in a symmetric similarity matrix for each marker. To constrain the analysis to spatially relevant regions, we applied a threshold of |z| > 3 to the ICN spatial maps from the Neuromark framework with functional 1.0 template^31^, thereby excluding weakly activated voxels and focusing on regions with strong functional engagement. Only voxels exceeding this threshold within each ICN were included in the similarity computation. Because the cell-type maps cover only cortical regions, cell-type microprofile networks were constructed using the 44 cortical ICNs, whereas neurotransmitter and mitochondrial microprofile networks were constructed using all 53 ICNs. This procedure yielded 49 region-by-region neurobiological microprofile networks: 24 for cell types, 19 for neurotransmitter receptors and transporters, and 6 for mitochondrial phenotypes, each capturing the similarity of biological feature distributions across functionally defined brain regions.

### Normative modeling construction

We constructed edge-level normative models for 1,378 FNC edges using a cohort of 3,834 neurotypical participants. Normative modeling was performed with the BLR^42^, as implemented in PCNtoolkit (version 0.30.post2; https://pcntoolkit.readthedocs.io/en/latest/)^43^, with cubic B-spline basis expansion to capture nonlinear age-related effects. For each FNC edge, connectivity strength was modeled as a function of age, sex, head motion, and site. Age-specific normative FNC matrices were then obtained by sampling the fitted normative model at 75 integer age points from 6 to 80 years. These matrices therefore represent model-estimated normative expectations. To quantify developmental and aging-related dynamics, we computed annualized FNC changes as first-order differences between normative FNC estimates at consecutive integer ages (e.g., FNC at age *t* minus FNC at age *t*–1), yielding 74 age-specific change patterns from 7 to 80 years. For the 1,515 individuals with psychiatric disorders, we derived individual d-scores for each FNC edge by comparing observed connectivity values against the model’s age-, sex-, motion-, and site-adjusted normative predictions. These d-scores reflect the magnitude and direction of deviation from expected normative connectivity at the single-subject level, enabling disorder-specific and cross-disorder characterization of FNC abnormalities.

### Linking normative FNC trajectories with neurobiological microprofile networks

To test whether normative FNC organization and its age-related changes are aligned with canonical neurobiological architecture, we quantified the edge-level spatial correspondence between model-estimated normative FNC patterns, annualized FNC-change patterns and neurobiological microprofile networks. For each age point, we constructed edge-level multiple linear regression models using the lower triangular elements of the FNC or FNC-change matrix as the dependent variable. Each model included the corresponding lower triangular elements of the neurobiological microprofile networks as predictors, ensuring that each observation corresponded to one FNC edge. Separate models were fitted for the three biological modalities: 24 cell-type, 19 neurotransmitter and 6 mitochondrial microprofile networks. For example, in the cell-type model at a given age, the expected FNC value of each edge was modeled as a linear combination of the 24 cell-type microprofile similarity values for that same edge. Analogous models were constructed for neurotransmitter and mitochondrial microprofile networks. The cell-type models were fitted using 946 unique edges among 44 cortical ICNs, whereas the neurotransmitter and mitochondrial models each included all 1,378 unique edges among 53 ICNs. Spatial confounds, including Euclidean distance and Dice similarity between ICNs, were regressed out prior to multivariate modeling. Model performance was quantified using adjusted *r*², which reflects the proportion of variance in normative FNC or FNC-change patterns explained by the combined neurobiological predictors. To evaluate the contribution of individual predictors, we conducted relative importance analysis^45,46^, which partitions the explained variance among predictors while accounting for correlations between them. Statistical significance was assessed using permutation-based null models. For each analysis, we generated 1,000 surrogate FNC maps by randomly permuting node positions, yielding null distributions for the adjusted *r*² and predictor-specific dominance. For normative FNC, *p*-values for adjusted *r*² were corrected across 75 age-points (6–80 years), whereas for age-related FNC changes, correction was performed across 74 age points (7–80 years). The *p*-values for predictors were corrected separately for each biological modality: 75 × 24, 75 × 19, and 75 × 6 comparisons for normative FNC, and 74 × 24, 74 × 19, and 74 × 6 comparisons for age-related FNC changes, corresponding to cell-type, neurotransmitter, and mitochondrial networks, respectively. Neurobiological markers that survived FDR correction and exhibited relative importance exceeding the average expected contribution (1/24 for cell-type, 1/19 for neurotransmitter, and 1/6 for mitochondrion) were identified as dominant individual predictors. To visualize lifespan patterns, we fitted locally estimated scatterplot smoothing (LOESS) curves to the empirical adjusted *r*² values across age, and compared them against LOESS curves fitted to selected percentiles (1st, 5th, 25th, 75th, 95th, 99th) of the null distributions.

### Disorder-specific and cross-disorder FNC alterations and their correspondence with neurobiological microprofile networks

We quantified both disorder-specific deviations and cross-disorder differences in FNC by analyzing edge-wise d-score maps across disorders. Disorder-specific alterations were identified using two-sided one-sample *t*-tests on d-scores for each disorder, whereas cross-disorder differences were assessed using two-sided two-sample *t*-tests for each pairwise comparison. All statistical tests were performed across 1,378 FNC edges, with FDR correction applied independently for each disorder and each pairwise disorder comparison. To assess whether these FNC alteration patterns were aligned with canonical neurobiological architecture, we constructed multivariate linear regression models for both disorder-specific and cross-disorder analyses. For each disorder, the disorder-specific Cohen’s *d* map served as the dependent variable; for each pairwise comparison, the cross-disorder Cohen’s *d* map was used instead. In all models, the neurobiological microprofile networks (i.e., cell-type, neurotransmitter, or mitochondrion) were entered as predictors. Spatial confounds (Euclidean distance and Dice similarity) were regressed out prior to modeling. Model performance was quantified using the adjusted *r*², reflecting the variance in disorder-related or cross-disorder FNC alterations explained by the neurobiological predictors. We assessed the contributions of individual markers using relative importance analysis^45,46^. Statistical significance was evaluated using one-sided permutation testing with 1,000 permutations to generate surrogate null models. For disorder-specific analyses, adjusted *r*² *p*-values were FDR-corrected across five disorder-level comparisons, and dominance-analysis *p*-values were corrected across modality-specific predictor counts (5 × 24 cell-type; 5 × 19 neurotransmitter; 5 × 6 mitochondrial). For cross-disorder analyses, adjusted *r*² *p*-values were corrected across 10 pairwise comparisons, and dominance-analysis *p*-values were corrected accordingly (10 × 24; 10 × 19; 10 × 6). Neurobiological markers that survived FDR correction and exhibited relative importance exceeding the average expected contribution (1/24 for cell-type, 1/19 for neurotransmitter, and 1/6 for mitochondrial features) were identified as dominant individual predictors.

### Donor-level reproducibility and lifespan stability of cell-type-specific microprofile networks

Because the present study links lifespan- and disorder-related FNC patterns to canonical microscale biological architecture, we evaluated whether the cell-type-specific microprofile networks used as reference maps were reproducible across transcriptomic donors and showed detectable age-related drift. We performed two complementary validation analyses using donor-level AHBA-derived cell-type maps and an independent lifespan transcriptomic dataset^44^. For the AHBA validation, donor-specific cell-type microprofile networks were constructed separately for each of the six AHBA donors. Donor-level reproducibility was assessed using a leave-one-donor-out strategy. For each donor and cell type, we averaged the corresponding networks from the remaining donors to obtain a leave-one-donor-out group-average reference network. We then computed the Spearman correlation between the donor-specific network and the leave-one-donor-out reference network. Statistical significance was assessed using permutation testing, and *p* values were corrected for 6 × 24 comparisons using the FDR procedure.

We next evaluated the lifespan stability of cell-type-specific regional architecture using the independent lifespan transcriptomic dataset^44^. Because this dataset does not provide sufficiently dense spatial sampling to construct microprofile networks analogous to those derived from the AHBA, we used region-pair contrast architecture as an approximation of cell-type-specific microprofile organization. To match the age range of the normative FNC analysis, we restricted the lifespan transcriptomic dataset to donors aged 6–80 years with available cortical data, yielding 19 donors with an observed age range of 8–70 years. Marker genes for the 24 cortical cell types used in the main analysis were mapped to the gene expression matrix. For each donor, cortical region and cell type, marker scores were computed by aggregating the expression of available marker genes^6^ and then averaging samples within each donor and cortical region. Regional marker-score profiles were rank-transformed within each donor and cell type, and all pairwise regional contrasts were computed as differences in rank-transformed marker scores between cortical region pairs. This yielded a donor-specific region-pair contrast vector for each cell type. Eleven common cortical regions were retained, resulting in 55 region-pair contrasts per cell type. Donor-level stability in the lifespan dataset was quantified using the same leave-one-donor-out framework. For each donor and cell type, the donor-specific region-pair contrast vector was correlated with the leave-one-donor-out group-average contrast vector using Spearman correlation. To test whether this stability measure changed with age, we assessed the association between donor-to-group contrast similarity and donor age for each cell type using Spearman correlation. Nonlinear age effects were further evaluated using cubic B-spline regression models, with adjusted *r*² used to summarize variance explained by nonlinear age terms. *P* values from age-association analyses were corrected across the 24 cell types using FDR correction.

## Data availability

The following neuroimaging datasets used in this study are available from public repositories or through controlled-access data-sharing platforms:

HCP-D (https://www.humanconnectome.org/study/hcp-lifespan-development),

HCP-YA (https://www.humanconnectome.org/study/hcp-young-adult),

HCP-A (https://www.humanconnectome.org/study/hcp-lifespan-aging),

LA5C (https://openfmri.org/dataset/ds000030/),

ABIDE I & II (https://fcon_1000.projects.nitrc.org/indi/abide/),

ADHD200 (http://fcon_1000.projects.nitrc.org/indi/adhd200/).

The FBIRN, COBRE, and CMDD datasets require application or controlled access and can be obtained from the respective project repositories or investigators.

Additional resources used in this study include:

Neuromark with functional 1.0 template (http://trendscenter.org/software/gift),

AHBA (http://human.brain-map.org/), snRNA-seq data

(https://cellxgene.cziscience.com/collections/d17249d2-0e6e-4500-abb8-e6c93fa1ac6f),

cell-type maps (https://trendscenter.org/data/),

lifespan transcriptomic dataset (https://hbatlas.org/pages/data),

neurotransmitter PET maps (https://github.com/netneurolab/hansen_receptors), and mitochondrial phenotype maps (https://identifiers.org/neurovault.collection:16418).

Study-generated data supporting the analyses presented in this work are publicly available at https://github.com/FelixFengCN/NeuroArchitecture-FNC^80^. Source data for Figs. 2–5 and Extended Data Figs. 1–10 are provided with this paper. Numerical source data for Supplementary Figs. 1 and 2 are provided as Supplementary Data.

## Code availability

Neuroimaging data processing and analysis were performed using FSL v6.0 (https://fsl.fmrib.ox.ac.uk/fsl/fslwiki/), SPM12 (http://www.fil.ion.ucl.ac.uk/spm/), and GIFT toolbox (http://trendscenter.org/software/gift; http://trendscenter.org/data) implemented in MATLAB R2024b. AHBA data were processed using the abagen toolbox (version 0.1.4; https://abagen.readthedocs.io/en/stable/) and cell-type imputation was performed using CIBERSORTx (https://cibersortx.stanford.edu/). Functional connectivity normative modeling was conducted using PCNtoolkit (version 0.30.post2; https://pcntoolkit.readthedocs.io/en/latest/). Relative importance analyses were performed using the relaimpo package (version 2.2.7) in R (version 4.5.0 development, 2025-02-15, revision 87726). Custom scripts supporting the downstream analyses are available at https://github.com/FelixFengCN/NeuroArchitecture-FNC. The version of the code associated with this study (v1.0.0) has been archived on Zenodo (https://doi.org/10.5281/zenodo.23573340)^80^.

## Acknowledgements

We thank all participants and investigators who contributed to the datasets used in this study. Data were provided in part by the Human Connectome Project (HCP), WU-Minn Consortium (Principal Investigators: David Van Essen and Kamil Ugurbil; 1U54MH091657), funded by the 16 NIH Institutes and Centers that support the NIH Blueprint for Neuroscience Research and by the McDonnell Center for Systems Neuroscience at Washington University. We also acknowledge the HCP-Development (HCP-D) and HCP-Aging (HCP-A) projects, supported by the National Institute of Mental Health (U01MH109589) and the National Institute on Aging (U01AG052564), respectively, and the McDonnell Center for Systems Neuroscience at Washington University.

We acknowledge the Function Biomedical Informatics Research Network (FBIRN; U24RR021992) and the Biomedical Informatics Research Network Coordinating Center (U24GM104203) for supporting access to the FBIRN data. We thank the Mind Research Network for sharing the COBRE dataset, whose data collection was supported by NIH grants 5P20RR021938/P20GM103472. Data from the UCLA Consortium for Neuropsychiatric Phenomics (LA5C) were obtained from the OpenfMRI database (ds000030). We thank the Autism Brain Imaging Data Exchange (ABIDE I and II) and ADHD-200 consortia for sharing their data. Primary support for ABIDE I was provided by NIMH grants K23MH087770 and R03MH096321, the Leon Levy Foundation, and gifts from Joseph P. Healy and the Stavros Niarchos Foundation to the Child Mind Institute. ABIDE II was primarily supported by NIMH grant R21MH107045.

We thank Jing Sui (Beijing Normal University) for providing the Chinese Han major depressive disorder (CMDD) dataset and for her methodological and analytical guidance, and Haojie Chen and Rongtao Jiang for their contributions to the interpretation of the findings.

We acknowledge the Allen Institute for Brain Science for providing the Allen Human Brain Atlas (AHBA), Jorstad and colleagues for sharing human cortical single-nucleus transcriptomic data through CELLxGENE, Hansen and colleagues and the original PET investigators for providing neurotransmitter maps, and Mosharov and colleagues for sharing mitochondrial phenotype maps through NeuroVault (collection 16418). We also thank Kang and colleagues for sharing the human brain transcriptomic dataset through the Human Brain Transcriptome (HBT) resource (GSE25219).

## Author contributions

G.F. and V.D.C. designed the research. Z.F. performed imaging data processing. G.F. processed all atlas data. V.D.C. provided methodological and analytical guidance. G.F. and V.D.C. developed the implementation code and software. J.C., A.I., and M.S.E.S. contributed to the interpretation of the findings. G.F. wrote the manuscript and prepared the figures. All authors reviewed and edited the manuscript. V.D.C. supervised and administered the project.

